# Single-cell RNA sequencing and large-scale bulk combination with machine learning reveal gastric cancer-related macrophage heterogeneity

**DOI:** 10.1101/2025.10.17.683046

**Authors:** Saisai Gong, Yu peng Zhao, Sheng Yang, Tong Liu, Cheng yun Li, Yun Zhou, Geyu Liang

## Abstract

**Background:** The tumor microenvironment (TME) significantly impacts cancer progression and overall patient survival. However, the complexity of tumor cell-TME interactions in gastric cancer (GC) and their underlying molecular basis remain to be systematically elucidated.

**Methods:** We performed single-cell RNA sequencing (scRNA-seq) on paired tumor and adjacent tissues from 3 treatment-naïve GC patients. We conducted an in-depth characterization of the cellular composition and molecular features of the GC TME, with a particular focus on tumor-associated macrophages (TAMs) subsets and their mediated intercellular communication. Investigating the effect of C1q on the polarisation of THP-1 induced macrophages through migration and invasion experiments

**Results:** Transcriptomic analysis of 56,151 single cells identified a key TAM subset – C1QA□ TAMs. This subset is characterized by high expression of genes including C1QA, C1QB, C1QC and FN1, exhibiting a distinct M2-like macrophage phenotype. We developed a LASSO-based predictive model, C1Q-LASSO, to accurately stratify patients based on survival outcomes and chemotherapy responses, independently of established prognostic parameters. In vitro functional experiments further confirmed that C1q directly promotes malignant phenotypes in GC cells. Additionally, C1q significantly enhances the malignant progression of gastric cancer induced by M2 macrophages.

**Conclusions:** Utilizing scRNA-seq technology, this study systematically delineated the heterogeneous landscape of the GC immune microenvironment and its complex cellular interaction network at single-cell resolution. We first identified and functionally validated the C1QA□ TAM subset, demonstrating its association with poor prognosis and pro-tumorigenic functions.

## 1. Introduction

Gastric cancer (GC) is a highly prevalent malignancy globally, ranking fourth in incidence and fifth in mortality worldwide^[1]^. In China, a significant proportion of GC patients are diagnosed at an advanced stage, resulting in poor survival outcomes, with a median overall survival often less than one year^[2, 3]^. Despite advancements in early diagnosis and treatment, the molecular mechanisms underlying GC initiation and progression remain incompletely elucidated. The 5-year survival rate for advanced-stage patients remains dismally low, below 5%^[4-6]^, underscoring the urgent need for deeper exploration of its biological basis and novel therapeutic targets.

The tumor microenvironment (TME), comprising immune cells (e.g., lymphocytes, myeloid cells), stromal cells, endothelial cells, and others^[7, 8]^, plays a central role in GC progression and treatment resistance. Among these components, tumor-associated macrophages (TAMs) are extensively implicated in poor patient prognosis and treatment resistance^[9, 10]^. TAMs significantly promote GC invasion and metastasis through diverse mechanisms, including mediating immunosuppression, remodeling the extracellular matrix, secreting pro-tumorigenic factors, and inducing metabolic reprogramming. Single-cell RNA sequencing (scRNA-seq) technology offers unprecedented resolution for dissecting the cellular and molecular heterogeneity of the TME, enabling the profiling of gene expression and metabolic characteristics at the single-cell level. Therefore, systematically characterizing the transcriptional profiles of key immune cells, particularly TAMs, within the GC TME is crucial for understanding the limitations of immunotherapy and developing novel TME-targeting therapeutic strategies to combat GC progression.

This study focuses on TAMs in the GC TME. We performed high-precision scRNA-seq on paired tumor and adjacent tissues obtained intraoperatively from three treatment-naïve GC patients. Through in-depth analysis of the TME’s cellular composition, we successfully identified a key TAM subset at the single-cell level: C1QA□ TAMs. This subset exhibits a characteristic M2-like macrophage phenotype accompanied by significant alterations in lipid metabolic activity. Importantly, integrating analysis with public transcriptomic datasets (TCGA-STAD) revealed that the enrichment of C1QA□ TAMs is significantly associated with poor prognosis in GC patients. Further in vitro functional experiments confirmed that C1q, a key molecule secreted by this subset, directly promotes malignant phenotypes in GC cells, including significantly enhanced proliferation, migration, and invasion capabilities.

Through a single-cell lens, this study uncovers a specific TAM subset (C1QA□ TAMs) with prognostic significance and pro-tumorigenic functions within the GC TME, along with its key effector molecule, C1q. These findings provide important experimental evidence and novel insights for deepening our understanding of the GC immune microenvironment, overcoming treatment resistance, and developing novel targeted therapeutic strategies.

## 2. Results

### 2.1 Single-cell sequencing identifies 11 major cellular subsets in GC

Fresh tumor tissues and adjacent non-tumor (paracancerous) tissues were collected from 3 treatment-naïve patients with non-metastatic GC. Following rapid enzymatic dissociation into single-cell suspensions, we performed single-cell transcriptome sequencing using droplet-based microfluidic technology (10x Genomics) (Fig. 1A). After stringent quality control, 56,151 high-quality cells were retained for analysis.

**Figure. 1.**
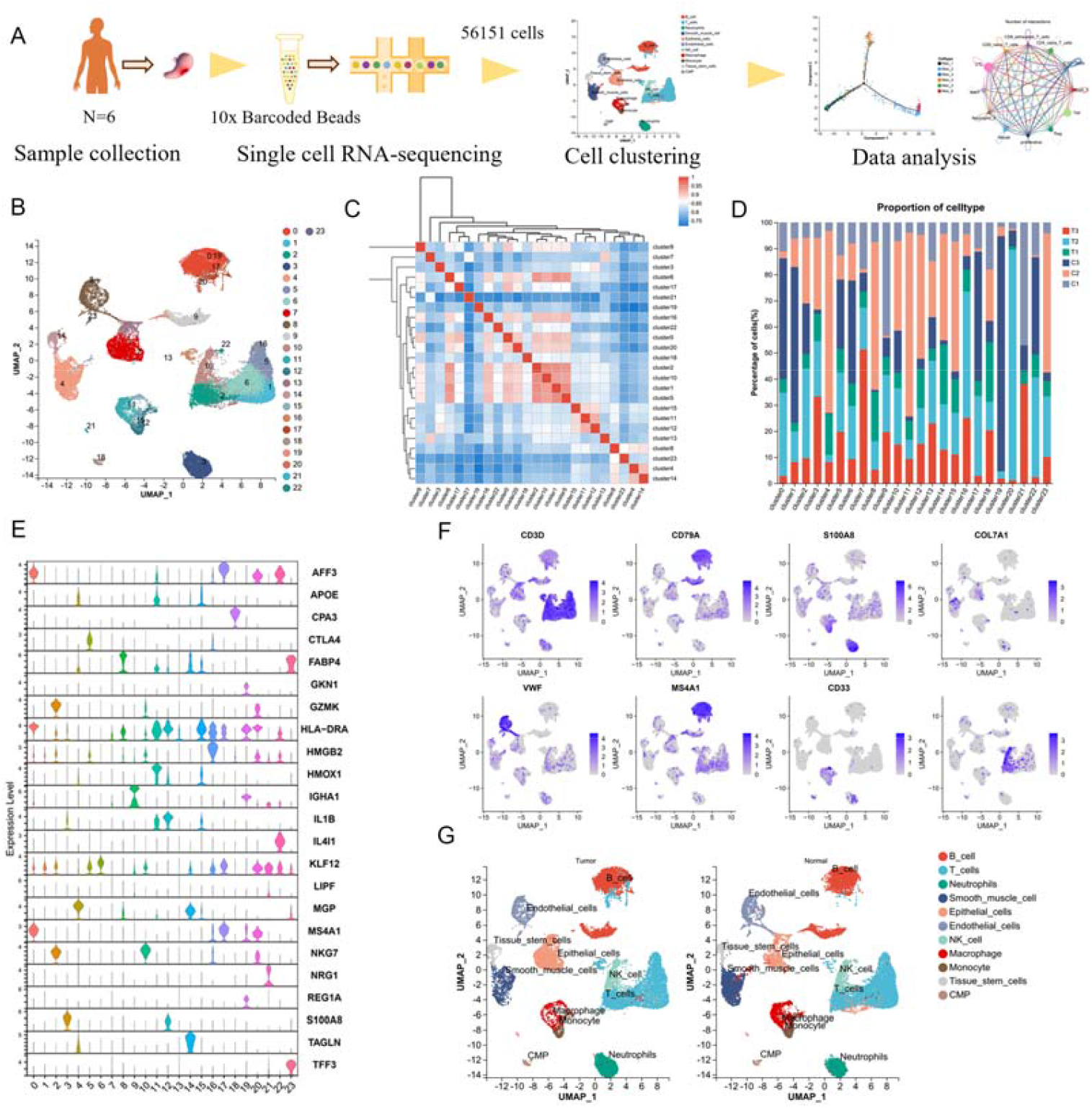
Single-cell atlas of gastric tumor and non-tumor gastric tissue. (A) Flowchart of single-cell sequencing data processing(n=3). (B) UMAP plots of 24 major cell types from GC and adjacent tissues, with 31,442 cells from adjacent tissues and 24,709 cells from GC tissues. (C) Heatmap of correlations between 24 cell subclusters. (D) The proportion of 24 cell subtypes in 3 samples. (E) Violin plot of the main marker expression distribution for 24 cell subtypes. (F) UMAP plot of the expression of classic marker genes for 24 cell subtypes. (G) UMAP shows 11 cell types of 56,151 cells.

Gene expression data underwent normalization and principal component analysis (PCA), followed by graph-based clustering which partitioned the cells into 24 transcriptionally distinct clusters (Fig. 1B). We performed inter-cluster correlation analysis based on top differentially expressed genes (DEGs) to assess cluster relationships (Fig. 1C). Cellular composition across clusters varied significantly between samples (Fig. 1D). Using canonical marker gene expression patterns (Fig. 1E-1F), we annotated and consolidated the 24 clusters into 11 major cell types (Fig. 1G), defined by their characteristic surface markers: B cells: CD79A, MS4A1; T cells: CD3D, CD3E; Natural killer (NK) cells: NCR1, NKG7; Epithelial cells: PGC, PGA3; Endothelial cells: RAMP2, PECAM1; Neutrophils: CSF3R, S100A8; Smooth muscle cells: ACTA2, TAGLN; Macrophages: MRC1, CD68; Monocytes: CD14, FCN1.

### 2.2 Heterogeneity of Myeloid Cells in the GC Microenvironment

Single-cell transcriptomic analysis identified 16 myeloid cell subpopulations (Fig. 2A), with macrophages (Clusters 0/1/5/6/8/11) characterized by high CD86/CD63/MRC1 expression, monocytes (Clusters 2/3/4) specifically expressing FCN1/S100A8, and dendritic cells (DCs; Clusters 7/9) showing low CD14 and high HLA-DR expression, while 5 clusters remained unassigned due to low quality or doublet status. Significant distribution differences existed between tumor and adjacent tissues (Fig. 2B). Monocyte analysis revealed CD14□/CD16□ populations formed two major pan-cancer clusters, indicating microenvironment-independent differentiation (Fig. 2C-2E). Although all monocytes expressed canonical markers (CD14/CD16/FCN1), tumor-infiltrating subsets upregulated HLA-DR/HLA-DRB1 and macrophage genes (CD86/CD163), confirming tissue-macrophage differentiation (Fig. S1A-S1B). Tumor-associated monocytes specifically overexpressed inflammatory factors (IL1B/CCL4/CXCL2/CXCR4), tissue-residency markers (NR4A1/NR4A2/NLRP3), growth factors (AREG/EGR1), and NF-κB pathway genes (NFKB1/NFKBIA). Among DCs, LAMP3□ cDC2 cells exhibited a migratory activated phenotype with upregulated migration/maturation regulators but downregulated TLR signaling (Fig. S1C-1D)—a conserved feature across cancers consistent with HCC reports^[11]^, indicating their role as mature DCs. Macrophages demonstrated significant heterogeneity^[12]^; the complement gene-high Mφ_4 subset (C1QA/C1QB/C1QC) displayed tissue-resident properties and pro-tumoral M2-like characteristics, with signature genes significantly enriched in TCGA pan-cancer data (Fig. 2F-2G).

**Figure. 2.**
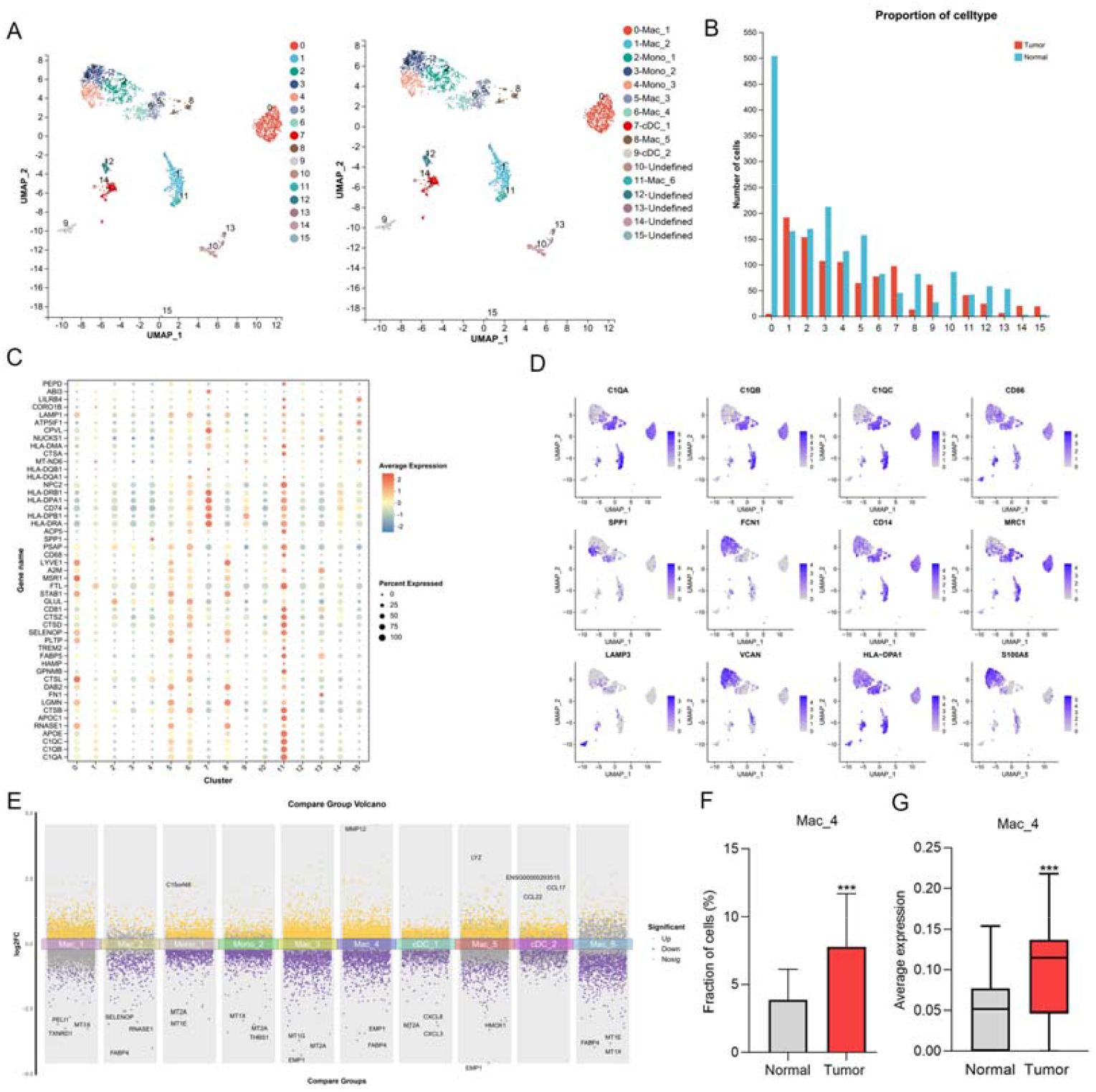
Single-cell sequencing characterization of tumor-infiltrating myeloid cells in GC. (A) UMAP of 16 myeloid cell subsets. (B) Proportion of myeloid cell subsets of tumors and adjacent tissues. (C) Characteristic gene expression bubble map of myeloid cell subsets. (D) Distribution of characteristic gene expression. (E) Differential gene analyses across subpopulations. (F-G) Levels of Mac_4 infiltration in TCGA STDA and expression levels of characterised genes.

### 2.3 Heterogeneous Transcriptional Features of Tumor-Associated Macrophages in GC

Re-clustering of GC macrophages identified six functionally distinct subsets: NRP1□ TAMs, FOS□ TAMs, MT-CO3□ TAMs, MRC1□ TAMs, HLA-DPA1□ TAMs, and C1QA□ TAMs (Fig. 3A). UMAP visualization revealed subset-specific gene expression patterns (Fig. 3B), wherein: C1QA□ TAMs exhibited high expression of M2-polarization markers and immunosuppressive genes (HAVCR2, SIRPA, LAIR1) with enhanced phagocytic activity; NRP1□ TAMs were enriched for angiogenesis-related genes (CD44, ITGAV); HLA-DPA1□ TAMs co-expressed M2 genes, immune checkpoints, the cancer stem cell core protein FN1 (which maintains proliferative and metastatic capacities)^[13]^, and immunosuppressive genes (SPP1, LAIR1, etc.) (Fig. 3C).

**Figure. 3.**
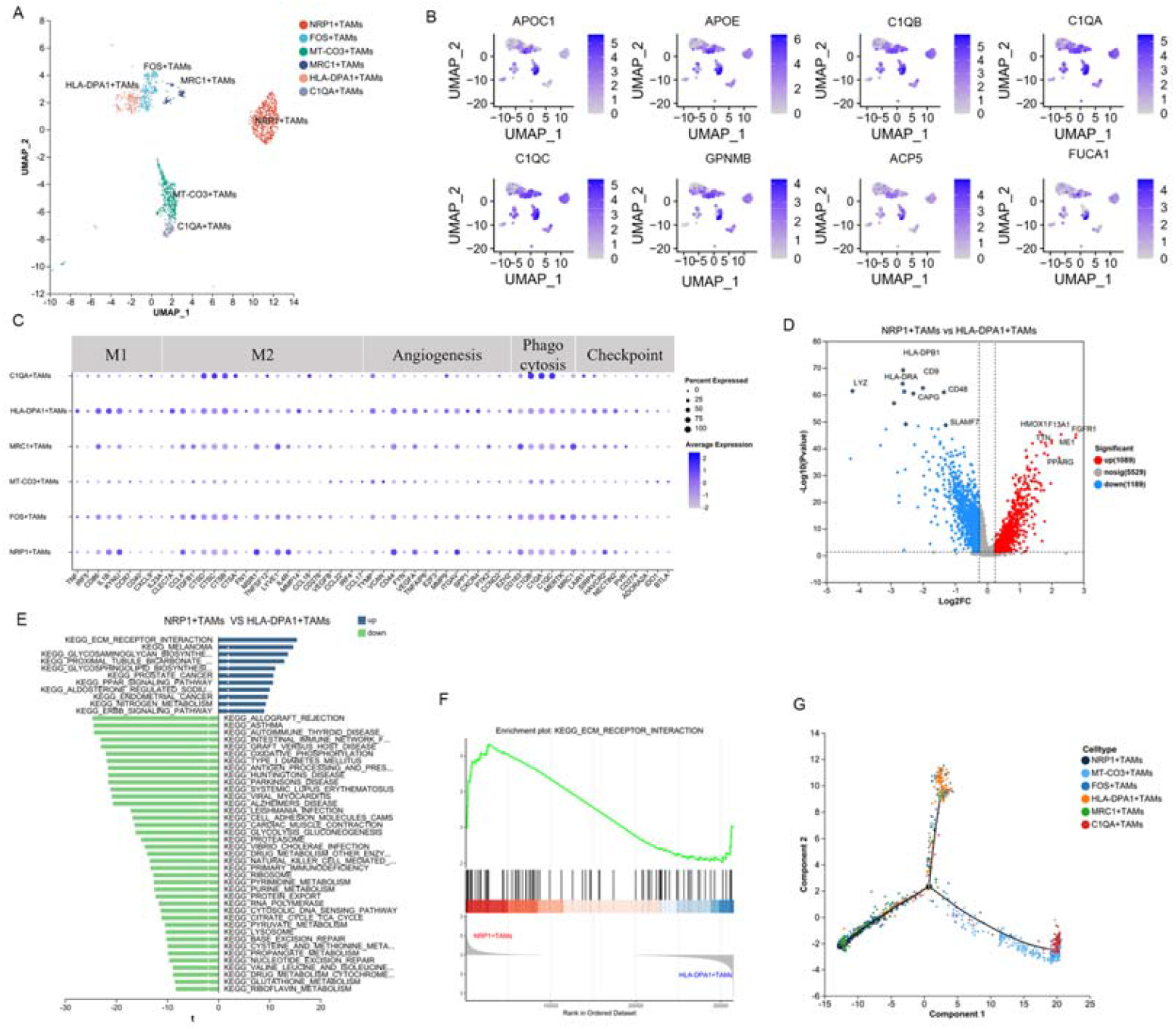
Molecular characterisation of tumour-associated macrophages in GC. (A) UMAP plot of T cells showing 14 clusters labelled with different colours. (B) UMAP dot of marker genes for different macrophage subpopulations. (C) Expression levels of M1-like, M2-like, angiogenic, phagocytic and checkpoint signature genes in macrophages. (D) Volcano maps of NRP1^+^ TAMs and HLA-DPA1^+^T AMs differential genes.(E) Gene set variation analysis (GSVA) identified the differentially expressed signaling pathways enriched in NRP1^+^ TAMs and HLA-DPA1^+^T AMs GSVA scores for differential genes.(F) Gene-set enrichment analysis was performed on gene sets of ECM receptor interaction. Positive NES indicate higher expression in NRP1^+^ TAMs.(G) Trajectory analysis of macrophage subpopulations.

Differential expression analysis (|log□FC| > 2, FDR < 0.05) showed significant upregulation of 1,089 genes in NRP1□ TAMs (Fig. 3D; Table S1). GSVA pathway enrichment demonstrated activation of ECM-receptor interaction and PPARG signaling in NRP1□ TAMs, alongside upregulated MHC class II antigen presentation in HLA-DPA1□ TAMs (Fig. 3E-F). Prognostic analysis using TCGA-STAD data confirmed that high expression of signature genes from both NRP1□ and HLA-DPA1□ TAMs predicted poorer survival (Fig. S2B). Pseudotime trajectory analysis suggested that NRP1□ and C1QA□ TAMs likely originate from MRC1□ TAMs (Fig. 3G), indicating macrophage reprogramming toward pro-tumorigenic phenotypes within the tumor microenvironment. This functional heterogeneity extends beyond classical M1/M2 classification paradigms.

### 2.4 Characterization of C1QA^+^ TAM-Mediated Cell Communication Networks

As found earlier, C1QA^+^TAMs were mainly expressed in M2 macrophages, and the presence of C1QA^+^TAMs was also found in other cancers, such as BRCA, KIRC, and NSCLC, based on scRNA-seq(Fig. S3A-S3B). The higher abundance of C1QA^+^TAMs in immuno-therapeutically treated KIRC suggests that the responsiveness of C1QA^+^TAMs in cancer immunotherapy is stronger. Analysis of ligand-receptor interactions using CellChat revealed significantly stronger intercellular communication within the GC microenvironment compared to adjacent normal tissues (Fig. 4A), evidenced by 2,370 interacting pairs detected in tumor tissues versus only 1,469 in paracancerous tissues (Fig. 4B-4C). Functioning as the central hub, C1QA^+^ TAMs established extensive connections with fibroblasts, stromal cells, endothelial cells, and immune cells—particularly Tregs and exhausted CD8□ T cells. Through ligand-receptor pairs including CXCL12-CXCR4, ANXA1-FPR1, and CCL3-CCR1 (Fig. 4D), they dynamically transmitted pro-tumorigenic signals.

**Figure. 4.**
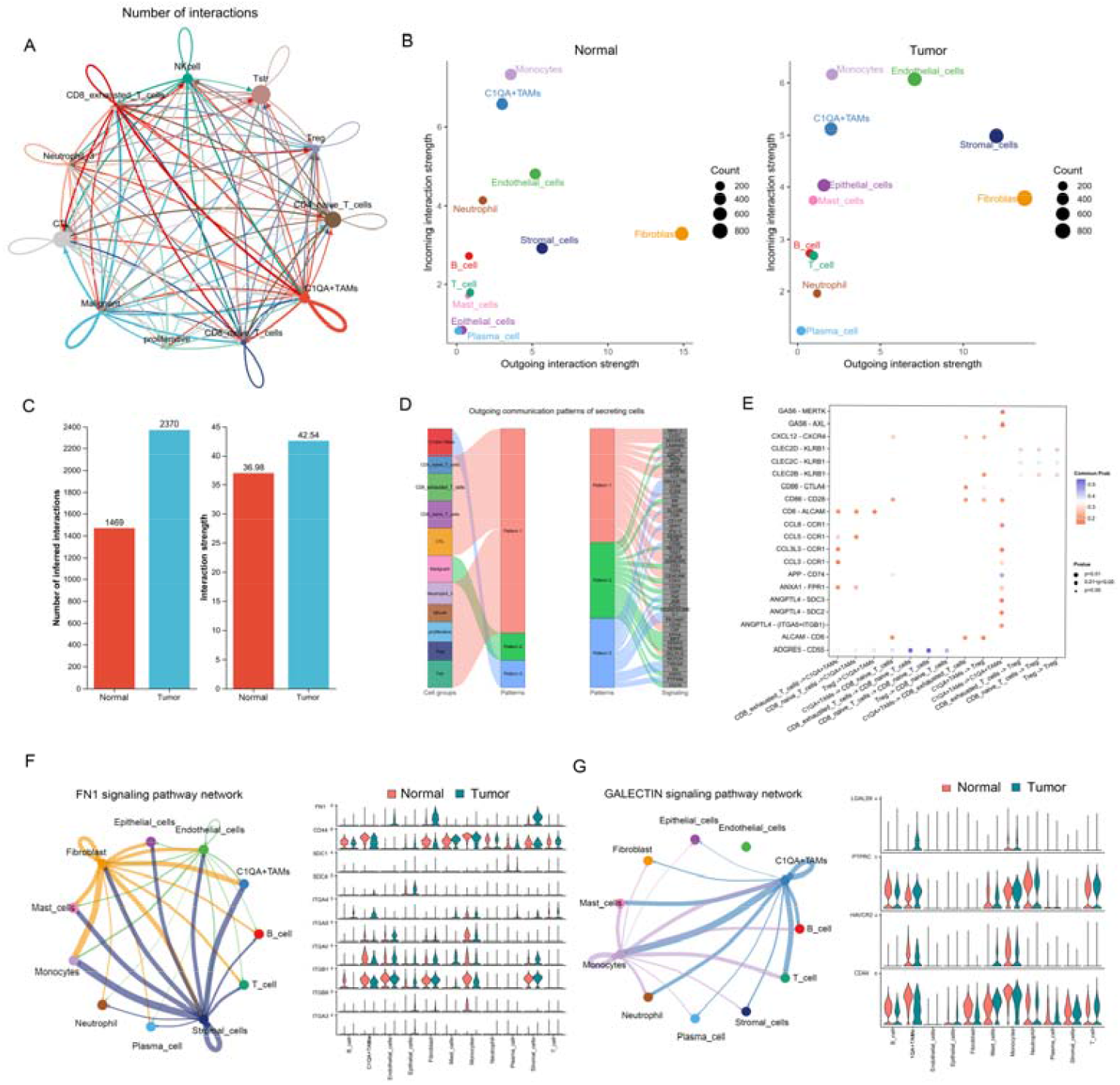
Inference of cellular communication from GC single-cell transcriptomes suggests a central role for C1QA^+^ TAMs. (A) Circle diagram showing the number of interations between various immune cells in the tumor. (B) Analysis of ligand-receptor numbers between tumor and normal tissue cells. (C) The bar graph represents the number of ligand-receptors in GC tumour and normal samples. (D) Demonstration of cellular communication patterns. (E) Dot plots show predicted ligand-receptor interactions between C1QA^+^ TAMs and various immune cells. (F-G) Analysis of ligand-receptor differences in the FN1 and SPP1 signalling pathways in tumour and normal samples.

Further communication pattern clustering analysis (Fig. 4E) demonstrated high activity of C1QA^+^ TAMs in pro-tumorigenic pathways such as SPP1, FN1, and GALECTIN signaling (Fig. 4F-4G). Notably, elevated SPP1 expression not only promoted tumor angiogenesis^1^ but also significantly correlated with poor patient prognosis (Fig. S4A-S4B). Collectively, this interaction network suggests that tumor cells recruit macrophages to collaboratively shape an immunosuppressive microenvironment.

### 2.5 Prognostic Role of C1QA^+^ TAMs in GC

To elucidate the functional role of C1QA^+^ tumor-associated macrophages (TAMs), we visualized the distribution of characteristic genes in GC using UMAP plots (Fig. 5A). The analysis revealed that C1QA^+^ TAMs exhibit high expression of complement genes (C1QA, C1QB, C1QC) and immunosuppressive genes (MRC1, NRP1, MARCO, TREM2) (Table S1). Furthermore, the abundance of C1QA^+^ TAMs was significantly higher in tumor tissues compared to adjacent normal tissues (Fig. 5B). KEGG pathway enrichment analysis indicated pro-tumorigenic characteristics of C1QA^+^ TAMs, showing enrichment in the VEGF signaling pathway and energy metabolism-related pathways, including oxidative phosphorylation and PPAR signaling pathway (Fig. 5C). This metabolic profile aligns with the known properties of M2 macrophages, which primarily rely on oxidative phosphorylation for energy production.

**Figure. 5.**
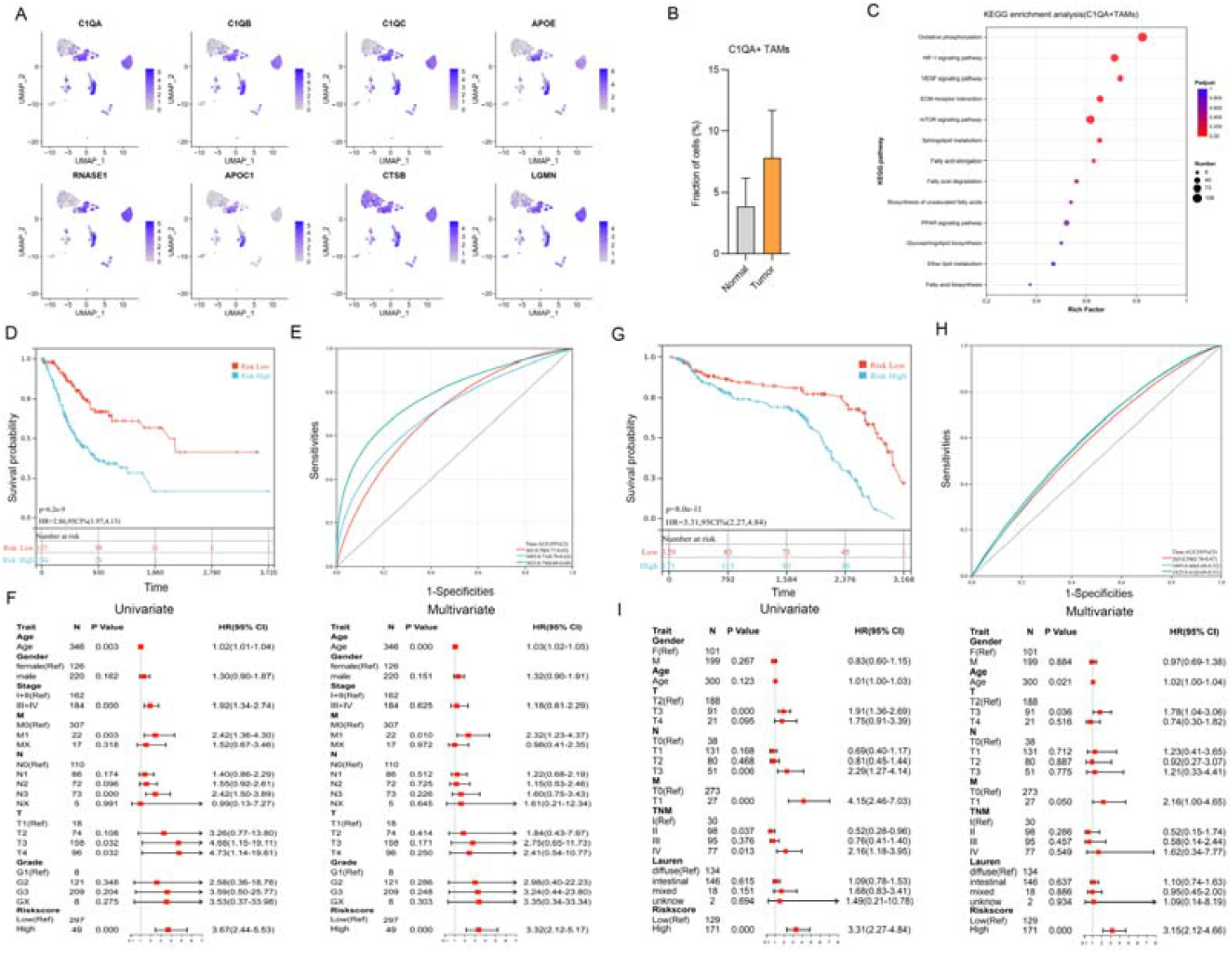
Impact of C1QA^+^TAMs signature genes in survival prognosis of GC. (A) Distribution of C1QA^+^TAMs signature gene expression in GC.(B) Abundance of C1QA^+^ TAMs in GC and paracancerous tissues.(C) KEGG pathway enrichment of highly expressed genes in C1QA^+^ TAMs.(D,E) Kaplan–Meier curve of TCGA and (G,H)GEO GC patients with high or low C1QA^+^ TAMs.(F) C1QA^+^ TAMs and clinical characteristics in TCGA and (I)GEO with univariate, multivariate cox risk analysis.

Analysis of C1QA^+^ TAMs signature gene expression in TCGA-STAD and GEO GC transcriptomic datasets confirmed that their abundance correlates with poorer overall survival in GC patients, consistent with previous reports. Concurrently, ROC curves based on this signature demonstrated good predictive efficacy for 1-year, 3-year, and 5-year survival rates (Fig. 5D-5H). Further univariate and multivariate Cox regression analysis identified both C1QA^+^ TAM abundance and tumor grade as independent risk factors for GC (Fig. 5F-5I). Thus, these findings collectively indicate that C1QA^+^ TAMs play a significant promotive role in the malignant progression of GC.

### 2.6 Development of a Subtype System Derived from C1QA+ TAMs for Predicting Patient Survival Prognosis

To assess the clinical efficacy of molecular typing of C1QA+TAMs in predicting patient survival outcomes, we combine bulk RNA-seq data, which are divided into a training set (TCGA, n=431), a validation set (GSE62254, n=300) and a test machine (GSE84437, n=351). Using the expression of C1QA+TAMs feature genes based on GLM, GBM, SVM, RF, KNN, NNET, DT, and Lasso Regression(Fig. 6A), we constructed predictive models, and after 10 repetitions of five-fold cross-validation to prevent overfitting, the lasso model showed the highest predictive accuracy, with an The AUC was 0.741 and this model was named C1Q-LASSO model(Fig. 6B).

**Figure. 6.**
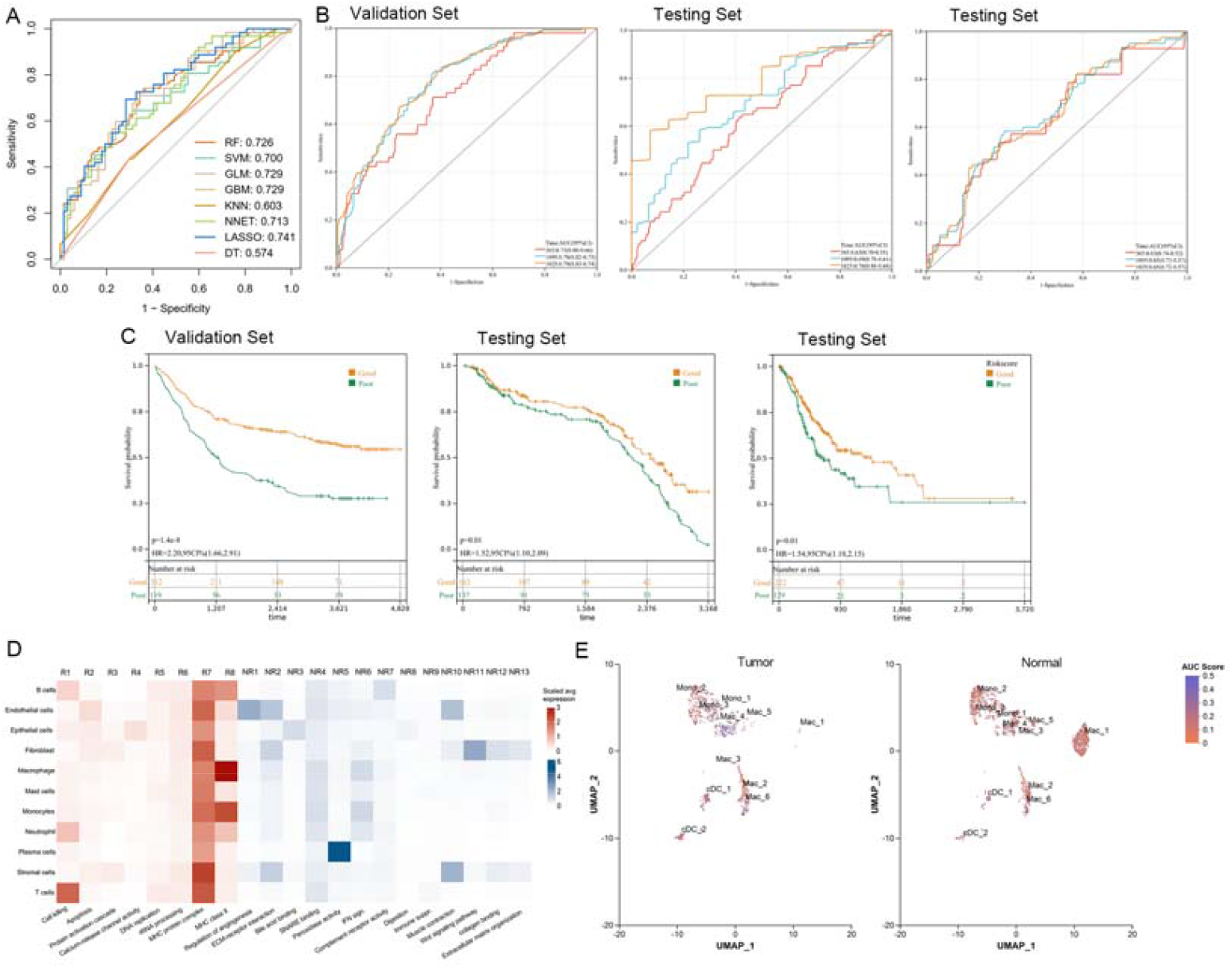
Prediction of survival prognosis by a model based on the characteristics of C1QA+TAMs and their responsiveness to immunotherapy. (A) Comparison of ROC curves for algorithmic performance of different machine learning models. (B) ROC curve representation of the applicability of the C1Q-LASSO model in the validation and test sets. (C) Kaplan-Meier curves comparing OS in both groups of patients in the validation and test sets. (D) Expression differential genes were clustered based on cell type in single-cell sequencing. (E) Annotation and UCell scoring of myeloid cells with gene cluster R8.

To further validate the stability and applicability of the C1Q-LASSO model, we tested it using independent data. This model demonstrated high predictive ability, and survival analysis also showed that patients with C1Q-LASSO subtype had lower survival time than patients with non-C1Q-LASSO subtype, which indicated its ability to stratify patients based on survival risk(Fig. 6C).

To assess the corresponding value of C1QA+TAMs in immune checkpoint therapy for gastric cancer, we obtained gene expression data from 45 gastric cancer patients treated with anti-PD-1. Differential gene expression analysis identified significantly high expression of 696 and 1382 genes in responding and non-responding patients, respectively. 8 gene clusters (R1-R8)were obtained by clustering 696 genes associated with the corresponding in single-cell data, and 13 gene clusters (NR1-NR13)were found by similarly clustering the set of genes with the non-corresponding(Fig. 6D). Genes associated with MHC II complexes were highly expressed in responding patients (R8), and C1QA, C1QB, and C1QC were highly expressed in Mac_6, suggesting a positive response of C1QA+TAMs in immunotherapy for gastric cancer(Fig. 6E).

### 2.7 C1q increased M2 macrophage-induced GC malignancy

To further validate the relationship between C1q and macrophage polarisation, we employed immunofluorescence to detect C1q and CD163 expression in gastric cancer patient samples. We observed that C1q was predominantly expressed in infiltrating CD163+ macrophages(Fig. 7A). Following C1q treatment, M0 macrophages exhibited increased expression of CD206, TGF-β, and IL-10(Fig. 7B). Co-culturing gastric cancer cells with conditioned medium from C1q-pretreated M0 macrophages revealed that C1q markedly promoted malignant cell migration(Fig. 7C). When IL-4 and IL-13 induced M2 macrophages were treated with C1q for 24 hours, C1q further enhanced the tumour-promoting capacity of M2 cells(Fig. 7D). These experiments demonstrate that C1q markedly promotes the polarisation of M2 macrophages.

**Figure. 7.**
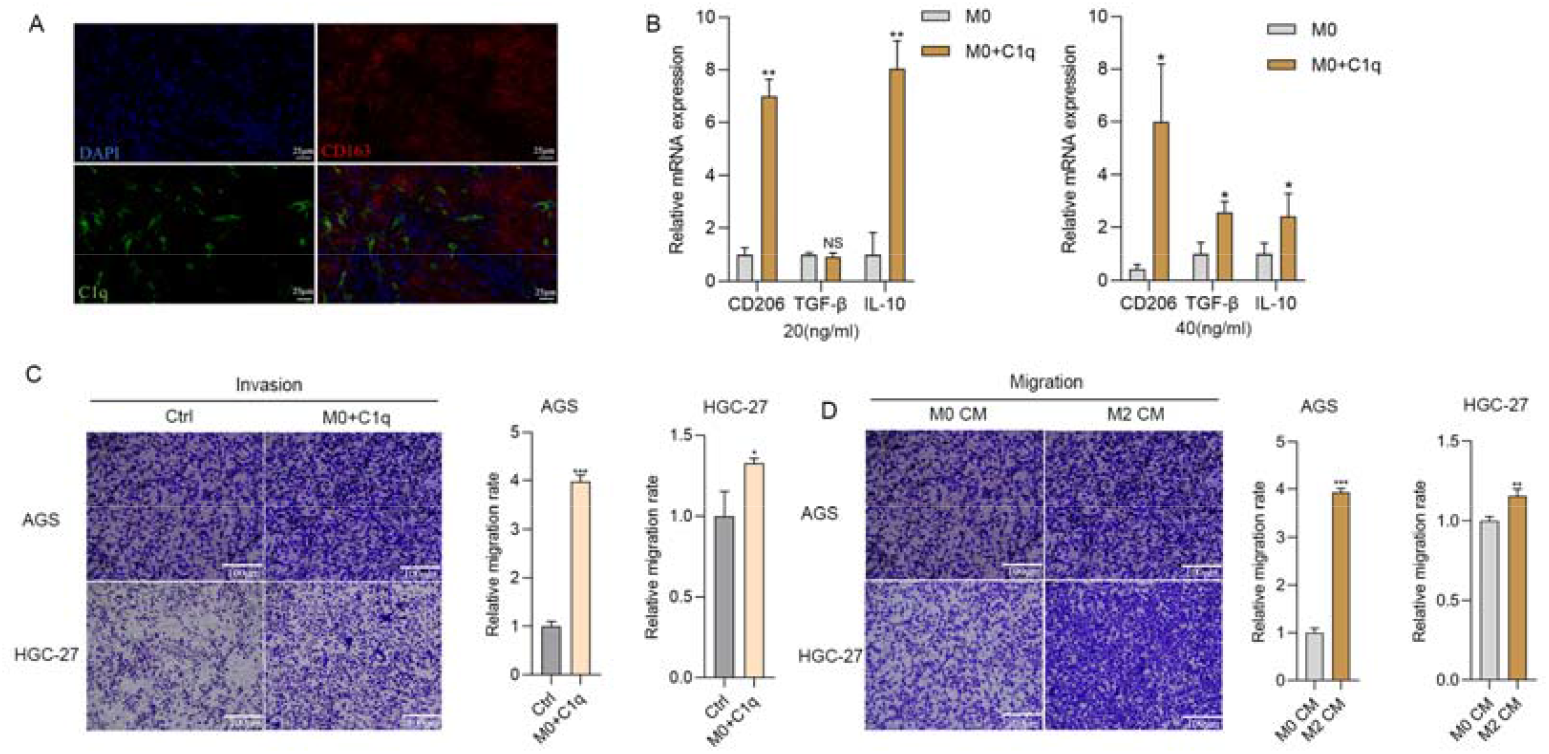
C1q is positively correlated with M2 macrophage polarisation in gastric cancer (A) Immunofluorescence staining for C1q and CD163 in gastric cancer tissue. (B) RT-qPCR validation of M2 macrophage marker expression levels. (C) Following treatment with M0 macrophage conditioned medium(CM), the invasion capacity of gastric cancer cells AGS and HGC-27 increased. (D) Following treatment with M2 macrophage CM, the migration capacity of GC cells increased.

### 2.8 C1q Promotes Malignant Progression in GC

Bioinformatics analysis revealed that high expression of C1QA^+^ TAMs is associated with a poorer prognosis in multiple cancers (Fig. 8A-B). To investigate the regulatory role of C1q in GC progression, GC cells (AGS and HGC-27) were treated with different concentrations of C1q. Results from CCK-8 assays (Fig. 8C) and cell scratching assay (Fig. 8D) demonstrated that C1q significantly promoted GC cell proliferation. Furthermore, transwell assays indicated that C1q enhanced both the migration (Fig. 8E) and invasion (Fig. 8F) of GC cells. These findings demonstrate that C1q facilitates malignant metastasis and acts as a promotive factor in GC progression.

**Figure. 8.**
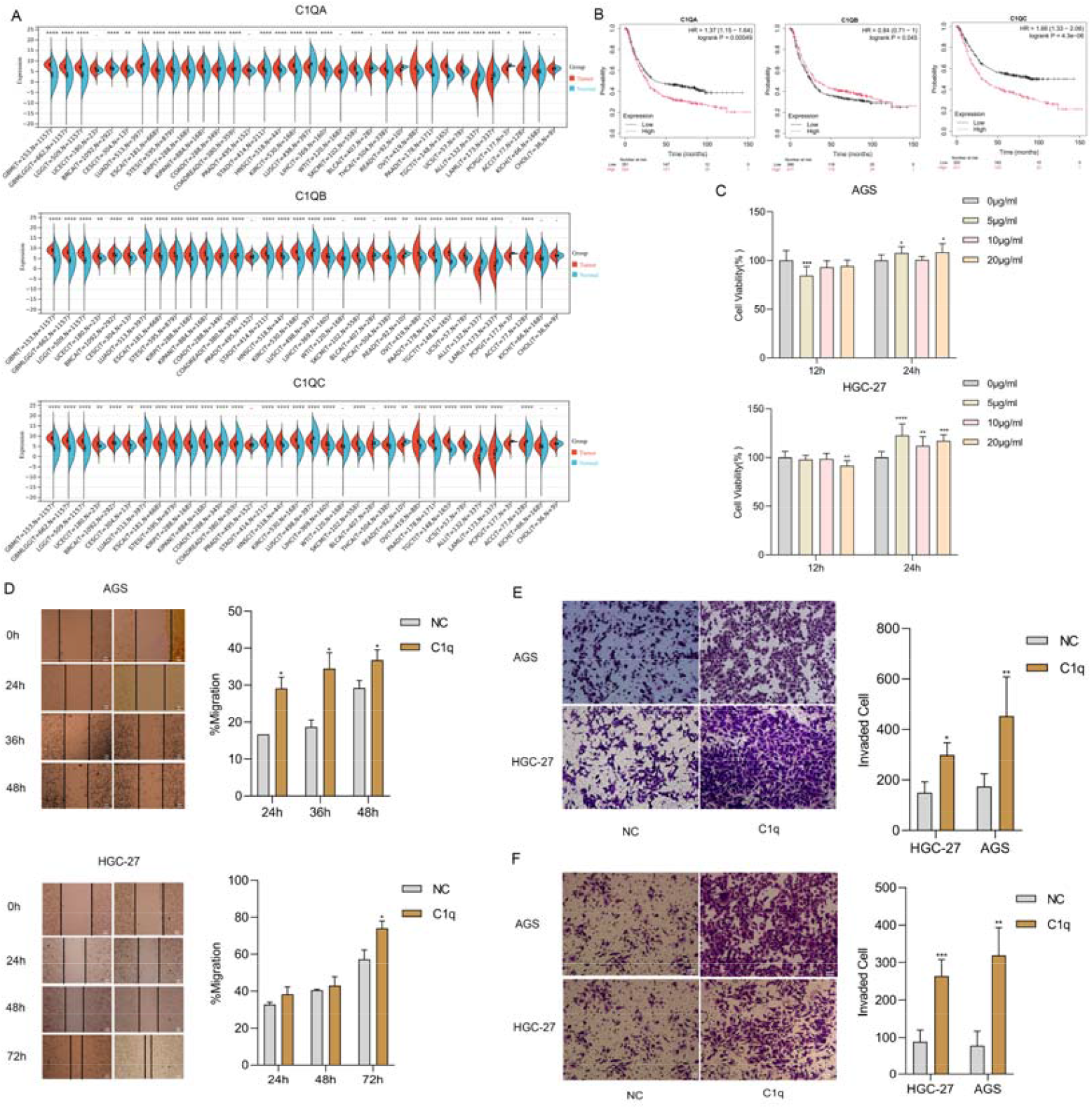
C1q Promotes Malignant Progression in GC (A)Differential expression of the C1Q gene in multiple cancers. (B) Survival curve analysis of C1q gene expression in GC. (C) CCK8 detects proliferation of GC cells. (D) Cell cloning experiments of GC cells. (E) C1q promotes migration and (F)invasion of GC cells.

## 3. Discussion

GC is a highly heterogeneous malignancy influenced by both genetic and environmental factors, posing significant challenges for diagnosis and treatment. Emerging single-cell technologies have become powerful tools for resolving tumor heterogeneity, enabling in-depth exploration of key biological processes including tumor progression stratification, immune tolerance, intercellular interactions, and tumor infiltration. These advances deepen our understanding of GC biology. This study comprehensively delineates the GC tumor landscape at single-cell resolution^[14-16]^ .

Our analysis revealed pronounced heterogeneity within the GC tumor microenvironment (TME), particularly among myeloid cell populations. We provided detailed characterization of conventional dendritic cell (cDC) heterogeneity. Dendritic cells (DCs) are classically categorized into conventional (cDCs) and plasmacytoid (pDCs) subtypes, with cDCs further differentiating into CD141□ and CD1C□ subsets^[17]^ .

LAMP3□ cDCs, recognized markers of DC maturation typically induced by CD40L stimulation, have been reported to originate from both cDCs and pDCs^[18-20]^ .

We identified and annotated a mature LAMP3□CCR7□ DC subset that elucidates myeloid cell functions in GC pathogenesis. These LAMP3□ DCs highly express chemokines (CCL19, CCL21) and co-stimulatory ligands (CCL17, CCL22), recruiting CCR4-expressing regulatory T cells (Tregs) to tumors and fostering an immunosuppressive microenvironment^[21, 22]^.Furthermore, tumor abundance of cDC-derived LAMP3□ DCs correlated significantly with infiltration levels of exhausted CD8□ T cells and Tregs^[11, 23]^ . Collectively, the LAMP3□ DC subset identified in GC mirrors populations observed across multiple cancers and likely participates in regulating T cell infiltration and functional exhaustion.

Tumor-associated macrophages (TAMs) constitute a key TME component^[24]^ . While historically simplified as pro-inflammatory (M1) or anti-inflammatory/pro-tumorigenic (M2) phenotypes based on murine models, this dichotomy inadequately captures TAMs complexity in human tumors^[25-27]^. Our scRNA-seq analysis identified six TAMs subsets predominantly enriched in tumor tissue. Among these, NRP1□ TAMs and C1QA□ TAMs exhibited stronger M2-like polarization. NRP1□ TAMs highly expressed pro-angiogenic signature genes, and their enrichment in tumor-promoting signaling pathways suggests roles in cancer progression and metastasis.

C1QA□ TAMs represent a tissue-resident macrophage subset^[28, 29]^ defined by core genes (C1Qs, APOE, TREM2, MRC1). C1q proteins enhance phagocytic function while suppressing pro-inflammatory cytokine production^[30-32]^, thereby exerting pro-tumor effects. We further discovered that C1QA□ TAMs exhibit distinct lipid metabolic signatures, with established links to maintaining immunosuppressive capacity through fatty acid metabolism regulation^[33, 34]^. Cell communication analysis revealed extensive interactions between C1QA□ TAMs and multiple exhausted CD8□ T cell/Treg populations, underscoring their role in shaping the immunosuppressive TME. Given predominant C1q expression in myeloid cells (especially macrophages), we confirmed through in vitro experiments that C1q protein significantly enhances GC cell proliferation and invasion. C1q significantly enhances the malignant progression of gastric cancer induced by M2 macrophages. This finding provides a mechanistic basis for the observed clinical association between high C1QA□ TAMs abundance and poor GC prognosis.

## 4. Methods

### 4.1 Patients and Samples

All patients participating in this study at Zhongda Hospital, Southeast University provided written informed consent. The study protocol was approved by the Ethics Committee of Zhongda Hospital, Southeast University. We enrolled three treatment-naïve GC patients whose diagnoses were pathologically confirmed. Patients had not received prior chemotherapy, radiotherapy, or other anticancer drug therapies. During surgery, fresh tumor tissue and matched paracancerous tissue were surgically resected. Paracancerous normal tissue was collected from regions >5 cm distal to the tumor margin to ensure its non-tumorous status. Detailed patient characteristics, including age and gender, are presented in Table S1.

### 4.2 Single-Cell Suspension Processing and Library Preparation

Single-cell suspensions passing quality control (viability >85%) were washed, resuspended to a concentration of 700–1200 cells/μL, and loaded onto the 10x Genomics Chromium™ system. This system generates Gel Beads-in-Emulsions (GEMs) to achieve single-cell partitioning. Cell barcoding was accomplished via reverse transcription (RT) of GEMs in a PCR thermal cycler. Subsequently, GEM emulsions were broken, and the barcoded first-strand cDNA was purified and enriched using magnetic beads. Following cDNA amplification and successful QC, Next-Generation Sequencing (NGS) libraries were constructed. Library preparation involved fragmentation, adapter ligation, and sample indexing via PCR. Final libraries underwent quantification and QC before being sequenced on either the Novaseq Xplus or DNBSEQ-T7 platform using a paired-end 150 bp (PE150) configuration. A targeted sequencing depth of ≥20,000 reads per cell was achieved.

### 4.3 Analysis of single cell sequencing data

Raw sequencing reads underwent initial quality assessment using Fastp, evaluating base composition (A/T/G/C) distribution and error rate profiles, followed by comprehensive processing through the 10x Genomics Cell Ranger pipeline which executed: STAR alignment to the reference genome; quantification of high-quality cells, genes per cell, and alignment rates; cell identification via barcode assignment; and gene expression quantification via UMI counting. To address inherent technical artifacts (empty droplets, multiplets, aberrant gene counts, elevated mitochondrial content), additional quality filtering was applied post-expression matrix generation using outlier removal criteria: genes per cell (nGene), UMIs per cell (nUMI), mitochondrial/ribosomal gene expression, and DoubletFinder-derived doublet detection, with systematic exclusion of cells exceeding ±2 standard deviations from parameter means to ensure downstream analytical robustness.

### 4.4 Dimension Reduction and Clustering

Following expression matrix filtration, Seurat implements default “LogNormalize” normalization: gene counts per cell are divided by total cellular transcripts (library size) to derive relative abundances, scaled by a factor of 10,000, and log-transformed. Post-normalization, technical variations are regressed out through linear modeling. Dimensionality reduction is performed using UMAP, while cell clustering identifies distinct subtypes based on shared transcriptional profiles. Resulting embeddings visually resolve cellular distributions.

### 4.5 Cell type and marker gene identification

Cell type annotation integrates marker gene analysis with computational tools (e.g., SingleR) through two primary approaches: (1) Supervised/semi-supervised marker-based identification leverages known cell-type-specific markers for manual/algorithmic annotation—straightforward yet susceptible to subjective bias and marker database limitations; (2) Unsupervised reference-based identification classifies cells by similarity to well-annotated expression datasets, minimizing human bias though constrained to broad categorizations due to reference scope limitations. To enhance reliability, these complementary methodologies are typically integrated.

### 4.6 Signature score analysis

To infer the functional status of macrophages subpopulations, we collected a list of gene signatures including M1, M2, angiogenesis, and phagocytosis-related gene signatures from a recent study by Cheng et al,. Signature scores for these gene signatures and pathways were calculated for each macrophage subpopulation using the ssGSEA method in the GSVA package.

### 4.7 Differential Gene Expression Analysis of Cellular Subpopulations

Following clustering analysis, cells were partitioned into distinct subclusters. Using Seurat, we performed comparative analysis between each subcluster and all remaining subclusters to identify differentially expressed genes (DEGs). The FindAllMarkers algorithm^[35]^ was employed to detect cluster-specific upregulated genes with the following thresholds: Expressed in ≥20% of cells within the cluster; Average log fold-change > 1.5; Adjusted p-value (FDR) < 0.05.

### 4.8 Gene Set Variation Analysis (GSVA)

We conducted GSVA^[36, 37]^using KEGG pathways and Gene Ontology gene sets from the Molecular Signatures Database (MSigDB v7.0). Pathway activity scores were calculated for each cell cluster using GSVA (v1.34.0), with results visualized in bar plots representing cluster-wise average expression levels.

### 4.9 Cell Communication Analysis

We inferred intercellular interactions between immune cells using CellChat (v0.0.1) ^[38]^. Interaction strengths were calculated based on expression levels of ligand-receptor pairs. Significantly enriched ligand-receptor pairs (P < 0.05) were extracted for further analysis.

### 4.10 Cell Trajectory Analysis

To determine potential linear differentiation between macrophage subpopulations, Monocle2^[39]^(version 2.26.0) was used and trajectories were inferred with cell subpopulation marker genes. A CellDataSet object was created using the normalised count data. Infer trajectories using default parameters after downscaling and cell sorting^[40]^.

### 4.11 TCGA data analysis

To validate the presence of C1QA^+^ TAMs, NRP1^+^ TAMs and HLA-DPA1^+^ TAMs in primary GAC in a large primary GAC cohort with available expression and clinical data, we performed ecotype deconvolution analyses using a bulk RNA sequencing dataset. We downloaded bulk RNA-seq data normalised to primary GC (STAD) generated by The Cancer Genome Atlas (TCGA).The RNA-seq data were processed and normalised by the TCGA using their transcriptome analysis pipeline for processing and normalisation.Clinical annotations of the TCGA-STAD cohort and molecular subtypes defined by the TCGA analysis working group were downloaded from the recent PanCanAtlas study. In addition, we downloaded GAC-seq data from the Gene Expression Omnibus database (GEO, https://www.ncbi.nlm.nih.gov/geo/). Raw gene expression values from microarray experiments were preprocessed (background corrected and log2 transformed) and quantile normalised using the Robust Multi-Array Averaging (RMA) algorithm.For datasets GSE84437, GSE62254,their original study-defined clinical, histopathological and survival data and molecular subtypes were downloaded and used for correlation analysis.

### 4.12 Cell proliferation assay (CCK-8)

Cells in the logarithmic growth phase were harvested, washed twice with phosphate-buffered saline (PBS), and dissociated using 0.25% trypsin solution. The cells were then resuspended in RPMI-1640 medium and gently pipetted to generate a single-cell suspension. After cell counting, cells were seeded into 96-well plates at a density of 8,000 cells per well. Cells were treated with 10 μg/mL C1q (Complement Technology) and incubated in a humidified 5% CO□ atmosphere at 37°C. Following treatment, 100 μL of serum-free medium containing 10% CCK-8 solution was added to each well. Plates were returned to the incubator for 2–4 hours. After incubation, the optical density (OD) at 450 nm was measured for each well using a microplate reader. Cell viability curves were generated based on the OD values.

### 4.13 Cell lines and culure

Human gastric cancer cells AGS and HGC-27 were purchased from the American Type Culture Collection (ATCC, USA). Cells were cultured in RPMI 1640 medium (Gibco, USA). THP-1 cells were induced to differentiate into M0 cells by treatment with 100 ng/mL phorbol 12-myristate 13-acetate (PMA; Sigma-Aldrich, USA) for 48 hours. M0 cells were polarised into M2 cells using 20 ng/mL IL-4 and IL-13.

### 4.14 Transwell migration assay

Cells in the logarithmic growth phase were harvested and seeded into the upper chamber of a transwell insert at a density of 1.5 × 10□ cells per well. Cells were treated with 10 μg/mL C1q. After 18 hours of incubation, the culture medium was aspirated and discarded. Cells remaining in the upper chamber were then: Washed three times with phosphate-buffered saline; Fixed with 4% paraformaldehyde for 10 minutes; Stained with crystal violet for 10 minutes. Finally, migrated cells on the lower membrane surface were counted and imaged.

### 4.15 Statistical analysis of data

Survival differences were calculated using the Kaplan-Meier method and compared by log-rank test. For data with non-normally distributed continuous variables, the Wilcoxon rank sum test was used for analysis. All data analyses were done using R software (v4.3.1) and all statistical tests were two-sided, with p < 0.05 considered statistically significant.

## Acknowledgments

The present study was supported by the National Natural Science Foundation of China (81972998), Postgraduate Research&Practice Innovation Program of Jiangsu Province(KYCX24_0483),Gansu Province Health Care Industry Research Program Projects (GSWSKY2024-09).

## Authors’ Contributions

Gongsaisai, Liang Geyu, Zhou Yun and Yang Sheng participated in the experimental design of the study. Liu Tong and Li Cheng yun conducted a statistical analysis. Gong Saisai and Zhao Yu peng contributed to the writing of the study protocol. All authors approved the final version of the manuscript.

## Supplementary Materials

Supplementary Table S1. Supplementary Figure S1,S2,S3,S4. (Please see supplementary materials).

## REFERENCES

1. Sung H, Ferlay J, Siegel RL, Laversanne M, Soerjomataram I, Jemal A, Bray F. Global Cancer Statistics 2020: GLOBOCAN Estimates of Incidence and Mortality Worldwide for 36 Cancers in 185 Countries. CA: a Cancer Journal For Clinicians 2021, 71(3): 209–249.

2. Ajani JA, Lee J, Sano T, Janjigian YY, Fan D, Song S. Gastric adenocarcinoma. Nature Reviews. Disease Primers 2017, 3: 17036.

3. Smyth EC, Nilsson M, Grabsch HI, van Grieken NC, Lordick F. Gastric cancer. Lancet (London, England) 2020, 396(10251): 635–648.

4. Galletti G, Zhang C, Gjyrezi A, Cleveland K, Zhang J, Powell S, Thakkar PV, Betel D, Shah MA, Giannakakou P. Microtubule Engagement with Taxane Is Altered in Taxane-Resistant Gastric Cancer. Clinical Cancer Research : an Official Journal of the American Association For Cancer Research 2020, 26(14): 3771–3783.

5. Hirata Y, Noorani A, Song S, Wang L, Ajani JA. Early stage gastric adenocarcinoma: clinical and molecular landscapes. Nature Reviews. Clinical Oncology 2023, 20(7): 453–469.

6. Zavros Y, Merchant JL. The immune microenvironment in gastric adenocarcinoma. Nature Reviews. Gastroenterology & Hepatology 2022, 19(7): 451–467.

7. Sahai E, Astsaturov I, Cukierman E, DeNardo DG, Egeblad M, Evans RM, Fearon D, Greten FR, Hingorani SR, Hunter T, Hynes RO, Jain RK, Janowitz T, Jorgensen C, Kimmelman AC, Kolonin MG, Maki RG, Powers RS, Puré E, Ramirez DC, Scherz-Shouval R, Sherman MH, Stewart S, Tlsty TD, Tuveson DA, Watt FM, Weaver V, Weeraratna AT, Werb Z. A framework for advancing our understanding of cancer-associated fibroblasts. Nature Reviews. Cancer 2020, 20(3): 174–186.

8. Gonzalez H, Hagerling C, Werb Z. Roles of the immune system in cancer: from tumor initiation to metastatic progression. Genes & Development 2018, 32(19-20): 1267–1284.

9. Busuttil RA, George J, Tothill RW, Ioculano K, Kowalczyk A, Mitchell C, Lade S, Tan P, Haviv I, Boussioutas A. A signature predicting poor prognosis in gastric and ovarian cancer represents a coordinated macrophage and stromal response. Clinical Cancer Research : an Official Journal of the American Association For Cancer Research 2014, 20(10): 2761–2772.

10. Wu Y, Grabsch H, Ivanova T, Tan IB, Murray J, Ooi CH, Wright AI, West NP, Hutchins GGA, Wu J, Lee M, Lee J, Koo JH, Yeoh KG, van Grieken N, Ylstra B, Rha SY, Ajani JA, Cheong JH, Noh SH, Lim KH, Boussioutas A, Lee J-S, Tan P. Comprehensive genomic meta-analysis identifies intra-tumoural stroma as a predictor of survival in patients with gastric cancer. Gut 2013, 62(8): 1100–1111.

11. Zhang Q, He Y, Luo N, Patel SJ, Han Y, Gao R, Modak M, Carotta S, Haslinger C, Kind D, Peet GW, Zhong G, Lu S, Zhu W, Mao Y, Xiao M, Bergmann M, Hu X, Kerkar SP, Vogt AB, Pflanz S, Liu K, Peng J, Ren X, Zhang Z. Landscape and Dynamics of Single Immune Cells in Hepatocellular Carcinoma. Cell 2019, 179(4).

12. Wu Y, Yang S, Ma J, Chen Z, Song G, Rao D, Cheng Y, Huang S, Liu Y, Jiang S, Liu J, Huang X, Wang X, Qiu S, Xu J, Xi R, Bai F, Zhou J, Fan J, Zhang X, Gao Q. Spatiotemporal Immune Landscape of Colorectal Cancer Liver Metastasis at Single-Cell Level. Cancer Discovery 2022, 12(1): 134–153.

13. Wang R, Song S, Qin J, Yoshimura K, Peng F, Chu Y, Li Y, Fan Y, Jin J, Dang M, Dai E, Pei G, Han G, Hao D, Li Y, Chatterjee D, Harada K, Pizzi MP, Scott AW, Tatlonghari G, Yan X, Xu Z, Hu C, Mo S, Shanbhag N, Lu Y, Sewastjanow-Silva M, Fouad Abdelhakeem AA, Peng G, Hanash SM, Calin GA, Yee C, Mazur P, Marsden AN, Futreal A, Wang Z, Cheng X, Ajani JA, Wang L. Evolution of immune and stromal cell states and ecotypes during gastric adenocarcinoma progression. Cancer Cell 2023, 41(8).

14. Savas P, Virassamy B, Ye C, Salim A, Mintoff CP, Caramia F, Salgado R, Byrne DJ, Teo ZL, Dushyanthen S, Byrne A, Wein L, Luen SJ, Poliness C, Nightingale SS, Skandarajah AS, Gyorki DE, Thornton CM, Beavis PA, Fox SB, Darcy PK, Speed TP, Mackay LK, Neeson PJ, Loi S. Single-cell profiling of breast cancer T cells reveals a tissue-resident memory subset associated with improved prognosis. Nature Medicine 2018, 24(7): 986–993.

15. Patel AP, Tirosh I, Trombetta JJ, Shalek AK, Gillespie SM, Wakimoto H, Cahill DP, Nahed BV, Curry WT, Martuza RL, Louis DN, Rozenblatt-Rosen O, Suvà ML, Regev A, Bernstein BE. Single-cell RNA-seq highlights intratumoral heterogeneity in primary glioblastoma. Science (New York, N.Y.) 2014, 344(6190): 1396–1401.

16. Bockerstett KA, Lewis SA, Wolf KJ, Noto CN, Jackson NM, Ford EL, Ahn T-H, DiPaolo RJ. Single-cell transcriptional analyses of spasmolytic polypeptide-expressing metaplasia arising from acute drug injury and chronic inflammation in the stomach. Gut 2020, 69(6): 1027–1038.

17. Binnewies M, Mujal AM, Pollack JL, Combes AJ, Hardison EA, Barry KC, Tsui J, Ruhland MK, Kersten K, Abushawish MA, Spasic M, Giurintano JP, Chan V, Daud AI, Ha P, Ye CJ, Roberts EW, Krummel MF. Unleashing Type-2 Dendritic Cells to Drive Protective Antitumor CD4^+^ T Cell Immunity. Cell 2019, 177(3).

18. Ishida T, Ueda R. CCR4 as a novel molecular target for immunotherapy of cancer. Cancer Science 2006, 97(11): 1139–1146.

19. Maier B, Leader AM, Chen ST, Tung N, Chang C, LeBerichel J, Chudnovskiy A, Maskey S, Walker L, Finnigan JP, Kirkling ME, Reizis B, Ghosh S, D’Amore NR, Bhardwaj N, Rothlin CV, Wolf A, Flores R, Marron T, Rahman AH, Kenigsberg E, Brown BD, Merad M. A conserved dendritic-cell regulatory program limits antitumour immunity. Nature 2020, 580(7802): 257–262.

20. Jin P, Han TH, Ren J, Saunders S, Wang E, Marincola FM, Stroncek DF. Molecular signatures of maturing dendritic cells: implications for testing the quality of dendritic cell therapies. Journal of Translational Medicine 2010, 8: 4.

21. Michea P, Noël F, Zakine E, Czerwinska U, Sirven P, Abouzid O, Goudot C, Scholer-Dahirel A, Vincent-Salomon A, Reyal F, Amigorena S, Guillot-Delost M, Segura E, Soumelis V. Adjustment of dendritic cells to the breast-cancer microenvironment is subset specific. Nature Immunology 2018, 19(8): 885–897.

22. Yoshie O, Matsushima K. CCR4 and its ligands: from bench to bedside. International Immunology 2015, 27(1): 11–20.

23. Zhang L, Li Z, Skrzypczynska KM, Fang Q, Zhang W, O’Brien SA, He Y, Wang L, Zhang Q, Kim A, Gao R, Orf J, Wang T, Sawant D, Kang J, Bhatt D, Lu D, Li C-M, Rapaport AS, Perez K, Ye Y, Wang S, Hu X, Ren X, Ouyang W, Shen Z, Egen JG, Zhang Z, Yu X. Single-Cell Analyses Inform Mechanisms of Myeloid-Targeted Therapies in Colon Cancer. Cell 2020, 181(2).

24. Ren X, Zhang L, Zhang Y, Li Z, Siemers N, Zhang Z. Insights Gained from Single-Cell Analysis of Immune Cells in the Tumor Microenvironment. Annual Review of Immunology 2021, 39: 583–609.

25. Mulder K, Patel AA, Kong WT, Piot C, Halitzki E, Dunsmore G, Khalilnezhad S, Irac SE, Dubuisson A, Chevrier M, Zhang XM, Tam JKC, Lim TKH, Wong RMM, Pai R, Khalil AIS, Chow PKH, Wu SZ, Al-Eryani G, Roden D, Swarbrick A, Chan JKY, Albani S, Derosa L, Zitvogel L, Sharma A, Chen J, Silvin A, Bertoletti A, Blériot C, Dutertre C-A, Ginhoux F. Cross-tissue single-cell landscape of human monocytes and macrophages in health and disease. Immunity 2021, 54(8).

26. DeNardo DG, Ruffell B. Macrophages as regulators of tumour immunity and immunotherapy. Nature Reviews. Immunology 2019, 19(6): 369–382.

27. Wang Z, Wu Z, Wang H, Feng R, Wang G, Li M, Wang S-Y, Chen X, Su Y, Wang J, Zhang W, Bao Y, Lan Z, Song Z, Wang Y, Luo X, Zhao L, Hou A, Tian S, Gao H, Miao W, Liu Y, Wang H, Yin C, Ji Z-L, Feng M, Liu H, Diao L, Amit I, Chen Y, Zeng Y, Ginhoux F, Wu X, Zhu Y, Li H. An immune cell atlas reveals the dynamics of human macrophage specification during prenatal development. Cell 2023, 186(20).

28. Wang J, Zhu N, Su X, Gao Y, Yang R. Novel tumor-associated macrophage populations and subpopulations by single cell RNA sequencing. Frontiers In Immunology 2023, 14: 1264774.

29. Sharma A, Seow JJW, Dutertre C-A, Pai R, Blériot C, Mishra A, Wong RMM, Singh GSN, Sudhagar S, Khalilnezhad S, Erdal S, Teo HM, Khalilnezhad A, Chakarov S, Lim TKH, Fui ACY, Chieh AKW, Chung CP, Bonney GK, Goh BK-P, Chan JKY, Chow PKH, Ginhoux F, DasGupta R. Onco-fetal Reprogramming of Endothelial Cells Drives Immunosuppressive Macrophages in Hepatocellular Carcinoma. Cell 2020, 183(2).

30. Lavin Y, Kobayashi S, Leader A, Amir E-AD, Elefant N, Bigenwald C, Remark R, Sweeney R, Becker CD, Levine JH, Meinhof K, Chow A, Kim-Shulze S, Wolf A, Medaglia C, Li H, Rytlewski JA, Emerson RO, Solovyov A, Greenbaum BD, Sanders C, Vignali M, Beasley MB, Flores R, Gnjatic S, Pe’er D, Rahman A, Amit I, Merad M. Innate Immune Landscape in Early Lung Adenocarcinoma by Paired Single-Cell Analyses. Cell 2017, 169(4).

31. Lazarov T, Juarez-Carreño S, Cox N, Geissmann F. Physiology and diseases of tissue-resident macrophages. Nature 2023, 618(7966): 698–707.

32. Braun DA, Street K, Burke KP, Cookmeyer DL, Denize T, Pedersen CB, Gohil SH, Schindler N, Pomerance L, Hirsch L, Bakouny Z, Hou Y, Forman J, Huang T, Li S, Cui A, Keskin DB, Steinharter J, Bouchard G, Sun M, Pimenta EM, Xu W, Mahoney KM, McGregor BA, Hirsch MS, Chang SL, Livak KJ, McDermott DF, Shukla SA, Olsen LR, Signoretti S, Sharpe AH, Irizarry RA, Choueiri TK, Wu CJ. Progressive immune dysfunction with advancing disease stage in renal cell carcinoma. Cancer Cell 2021, 39(5).

33. Ling GS, Crawford G, Buang N, Bartok I, Tian K, Thielens NM, Bally I, Harker JA, Ashton-Rickardt PG, Rutschmann S, Strid J, Botto M. C1q restrains autoimmunity and viral infection by regulating CD8^+^ T cell metabolism. Science (New York, N.Y.) 2018, 360(6388): 558–563.

34. Zhang S, Peng W, Wang H, Xiang X, Ye L, Wei X, Wang Z, Xue Q, Chen L, Su Y, Zhou Q. C1q^+^ tumor-associated macrophages contribute to immunosuppression through fatty acid metabolic reprogramming in malignant pleural effusion. Journal For Immunotherapy of Cancer 2023, 11(8).

35. Butler A, Hoffman P, Smibert P, Papalexi E, Satija R. Integrating single-cell transcriptomic data across different conditions, technologies, and species. Nature Biotechnology 2018, 36(5): 411–420.

36. Hänzelmann S, Castelo R, Guinney J. GSVA: gene set variation analysis for microarray and RNA-seq data. BMC Bioinformatics 2013, 14: 7.

37. Liberzon A, Subramanian A, Pinchback R, Thorvaldsdóttir H, Tamayo P, Mesirov JP. Molecular signatures database (MSigDB) 3.0. Bioinformatics (Oxford, England) 2011, 27(12): 1739–1740.

38. Jin S, Guerrero-Juarez CF, Zhang L, Chang I, Ramos R, Kuan C-H, Myung P, Plikus MV, Nie Q. Inference and analysis of cell-cell communication using CellChat. Nature Communications 2021, 12(1): 1088.

39. Cao J, Spielmann M, Qiu X, Huang X, Ibrahim DM, Hill AJ, Zhang F, Mundlos S, Christiansen L, Steemers FJ, Trapnell C, Shendure J. The single-cell transcriptional landscape of mammalian organogenesis. Nature 2019, 566(7745): 496–502.

40. Street K, Risso D, Fletcher RB, Das D, Ngai J, Yosef N, Purdom E, Dudoit S. Slingshot: cell lineage and pseudotime inference for single-cell transcriptomics. BMC Genomics 2018, 19(1): 477.

